# Notes on the endemic and near endemic threatened rainforest plant species of Banyang Mbo Wildlife Sanctuary, SW Region, Cameroon, and *Tricalysia banyangmbo* sp nov. (Rubiaceae-Coffeeae) a new coffee relative

**DOI:** 10.1101/2025.03.03.616975

**Authors:** Martin Cheek, Bonaventure Sonké

## Abstract

Species new to science discovered from the Banyang Mbo Wildlife Sanctuary (BMWS) of SW Region Cameroon are reviewed and summarised, as are species listed as threatened which have been recorded, both from A. lowland evergreen forest and B. from cloud (submontane) forest. In total five globally Critically Endangered, 17 Endangered and 21 Vulnerable plant species are listed for BMWS.

We describe an additional narrow endemic of BMWF, *Tricalysia banyangmbo sp. nov*. (Sect. *Tricalysia* Rubiaceae-Coffeeae), similar in the subulate calyx lobes exceeding the calyx tube, and in the stems being densely patent shortly hairy to *T. sylvae* Robbr. of Littoral, South Region, Cameroon and Gabon. The new species differs in having stipules in which the awn is shorter than the blade (vs vice versa), the leaf bases are rounded (vs cordate), the blades lack domatia (vs present) and the flowers are 4([ndash] 5)-merous (vs 5 &[ndash] 6&[prime] merous), the anthers lacking a distinct and conspicuous apical connective appendage (vs present). The new species is known from three collection sites with an estimated area of occupation of c. 4 km^2^ and is in an area threatened by oil palm plantations. We provisionally assess the species with the 2012 IUCN standard as Critically Endangered CR B1ab(iii) + B2ab(iii).

## Introduction

### The genus *Tricalysia*

The new species was readily placed in *Tricalysia* of tribe Coffeeae (Rubiaceae) by the combination of shortly sheathing interpetiolar stipules with awns, contracted axillary inflorescences with a series of 3(–4) cupular calyculi subtending the flowers, calyx teeth and tube conspicuous, corolla tubes about as long as lobes, anthers largely exserted, inserted at the mouth of the corolla tube which lacks exserted hairs. Five sections were hitherto recognized in *Tricalysia* s.str. (Robbrecht 1982, 1983, 1987; Ranarivelo-Randriamboavonjy et al. 2007) but this classification is clearly in need of a revision. According to Lachenaud et al. (2020), most of these sections were based on few key characters: unisexual flowers in sect. *Androgyne* Robbr. which includes the Malagasy species, free bracteoles in sect. *Probletostemon* (K.Schum.) Robbr., anthers fully or partly included in sect. *Ephedranthera* Robbr., and calyx longitudinally split in sect. *Rosea* (Klotzsch) Robbr.; the last section, sect. *Tricalysia*, being essentially defined by the lack of all these features. In fact, some of these characters are not always reliable. A molecular study of the group (Tosh et al. 2009) suggested that most of the sections are not monophyletic, but the resolution and sampling were not sufficient to propose an alternative classification.

The genus *Tricalysia* currently contains about 79 described species, which are mainly evergreen hermaphroditic shrubs or small trees of lowland evergreen forest in tropical Africa from Senegal and Guinea in the west (Gosline et al. 2023) to Sudan in the N and E (Darbyshire et al. 2015) extending to South Africa (but absent from Namibia and Botswana) and to Madagascar (Robbrecht 1987a). However, several species are subshrubs of drier habitats in southern Africa, and at least one species is deciduous. Formerly, the name *Tricalysia* was extended to Asian species now included in *Diplospora* DC. and *Discospermum* Dalzell (Ali & Robbrecht 1991). Those Asian species are distinct from *Tricalysia* s.s. (in Africa) by their unisexual tetramerous flowers. *Tricalysia* is now restricted to Africa and Madagascar. *Sericanthe* Robbr., an African genus segregated from *Tricalysia* by Robbrecht (1978), is distinguished by seeds with a scar-like area from apex to ventral face, basifixed anthers, often possessing bacterial nodules in the leaves and a sericeous indumentum on the outer corolla surface (Bridson & Verdcourt 2003). The molecular phylogenetic analysis of *Tricalysia* by Tosh et al. (2009) showed that the former *Tricalysia* subgenus *Empogona* had a sister relationship with *Diplospora*. Accordingly, the African genus *Empogona* Hook.f. was resurrected. Its species are distinguished from *Tricalysia* in the strict sense by black fruits, flag-like anther connectives, hairs projecting from the corolla throat and free, often alternate distal bracts. The phylogeny of the group was refined by Arriola et al. (2018) who included Asian representatives in the sampling and postulated that *Tricalysia sensu stricto* is sister to a clade (*Sericanthe* (*Diplospora, Empogona*)). A key to the 12 genera in tribe *Coffeeae* can be found in Cheek et al. (2018a). This tribe has its highest generic diversity in Cameroon with nine of the genera present.

Most African *Tricalysia* species are well studied as a result of Robbrecht’s revisions (Robbrecht 1979, 1982, 1983, 1987a, 1987b). Within *Tricalysia* as currently defined, Robbrecht (1979, 1982, 1983, 1987a) recognised five sections in Africa: Sect. *Ephedrantha* Robbr., Sect. *Probletostemon* (K.Schum.) Robbr., Sect. *Tricalysia*, Sect. *Rosea* (Klotzsch) Robbr. and an unnamed Madagascan section. However, only eleven species of the 78 accepted *Tricalysia* were sampled in the phylogenetic study by Tosh et al. (2009), and of these, only six were African, so further, more intensively sampled, phylogenetic studies are called for if the current sectional classification is to be tested. Since the monographic studies of Robbrecht, seven new taxa have been published from the Flora Zambesiaca region by Bridson (in Bridson & Verdcourt 2003) and nine new taxa published for the Madagascan section now named Sect. *Androgyne* Robbr. (Ranarivelo-Randriambovanjy et al. 2007). Apart from these, only six other new species have been published, three from Cameroon: *Tricalysia lejolyana* Sonké & Cheek (Sonké et al. 2002a), *T. achoundongiana* Robbr., Sonké, & Kenfack (Sonké et al. 2002b) and *T. elmar* Cheek (Cheek et al. 2020); two from Gabon and one from Equatorial Guinea (Lachenaud et al. 2020). However, further new species are likely to be discovered in Gabon, where 118 *Tricalysia* specimens are recorded as being unidentified to species versus 219 identified (Sosef et al. 2006: 373 – 375), and in Cameroon, where six unidentified, possibly undescribed species are recorded from the Bakossi area alone in Cheek et al. (2004). As Robbrecht (1987a) stated of “a number of inadequate specimens that probably represent several more species.”: “The rain forests of esp. Cameroun and Gabon are undoubtedly underexplored areas where more material of these and other suspected new taxa may be discovered”

Many species of the genus are geographically localised, rare and threatened. For example, of the 29 taxa listed for Cameroon (Onana 2011), twelve species of *Tricalysia* were assessed as threatened (Onana & Cheek 2011) mainly because they are known from a single or few locations, and some are threatened by logging followed by agriculture such as palm oil plantations, e.g. *Tricalysia lejolyana* (Sonké et al. 2002a).

Within *Tricalysia*, the new species keys out to Sect. *Tricalysia* because the calyx tube is well developed, the calyx lobes are subulate and the corolla throat is glabrous. In Robbrecht’s (1987) key to species, the material keys out to *Tricalysia sylvae* Robbr. because the calyx has a single split, the calyx lobes are subulate, domatia are absent, the inflorescences are condensed and the young stems and petioles are densely covered in erect hairs. However, the material, while similar in many features to this species, differs in many qualitative and quantitative characters as well as being geographically separated (see Table 1) and is accordingly here formally described and named as *Tricalysia banyangmbo* Cheek & Sonké.

**Table 1.** Diagnostic characters separating *Tricalysia sylvae* from *Tricalysia banyangmbo*. Data on *Tricalysia sylvae* from Robbrecht (1987a) and specimens at K.

|  | *Tricalysia sylvae* | *Tricalysia banyangmbo* |
| --- | --- | --- |
| Indumentum of stem (length, mm) | 0.5(– 1) | 0.2 – 0.25 |
| Leaf-blade shape & maximal dimensions (cm) | Obovate 14 x 6 | Elliptic or elliptic-oblong, less usually oblanceolate 10.1 x 3.9 |
| Lateral nerve number (each side of midrib) | 4 – 7 | (3 – )4(– 5) |
| Leaf base shape | Cordate | Rounded |
| Domatia | Present | Absent |
| Stipule blade shape/awn length (mm) | Triangular/2 – 5 | Hemi-orbicular/0.75 – 1 |
| Flowers | 5 – 6-merous | 4(– 5)-merous |
| Corolla lobe dimensions | 3.5 – 4.5 x 1.2 – 2 | 4 – 7 x 1 – 1.5 |
| Anthers | Medifixed | Submedifixed |
| Anther apical connective | Present | Absent |
| Geographic range | Littoral Region, Cameroon to Gabon | SW Region, Cameroon |

The new species of *Tricalysia* A.Rich. reported in this paper was discovered as a result of the long-term survey of plants in Cameroon to support improved conservation management led by botanists from Royal Botanic Gardens, Kew, the IRAD (Institute for Research in Agronomic Development)– National Herbarium of Cameroon, Yaoundé and University of Yaoundé I. This study has initially focussed on the Cross-Sanaga interval (Cheek et al. 2001) which contains the area with the highest species and generic diversity per degree square in tropical Africa (Barthlott et al. 1996; Dagallier et al. 2020). To date, three new genera have been discovered through these studies, *Kupea* Cheek & S.A. Williams (Cheek et al. 2003), *Korupodendron* Litt & Cheek (2002) and *Kupeantha* Cheek (Cheek et al. 2018) and hundreds of new species (including Alvarez-Aguirre et al. (2021), Champluvier & Darbyshire (2009), Cheek (2000, 2003, 2017), Cheek & Ameka (2008), Cheek & Csiba (2000, 2002a,b), Cheek & Etuge (2009a,b,c), Cheek & Fischer (1999), Cheek & Ngolan (2007), Cheek & Onana (2021), Cheek & Sonké (2005), Cheek & Xanthos (2012), Cheek et al. (2002a, b, 2008, 2009, 2017, 2018b, 2020a, 2021, 2022, 2023), Couvreur et al. (20220, Darbyshire et al. (2011), Ghogue et al. (2017), Gosline & Cheek (1998), Gosline et al. (2014, 2022), Hoffmann & Cheek (2003), Kenfack et al. (2003), Lachenaud (2019), Lachenaud et al. (2013, 2024), Maas-van de Kamer et al. (2016), Mackinder et al. (2010), Mackinder & Pennington (2011), Muasya et al. (2010), Onana & Chevillotte (2015), Prance & Jongkind (2015), Sonké (2000), Sonké et al. (2005, 2006, 2007, 2008), (Stoffelen et al. 1997), (Stone et al. 2008) van der Burgt et al. (2015)).

The herbarium specimens collected in these surveys formed the primary data for the series of Conservation Checklists that began at Mt Cameroon (Cheek et al. 1996), with the Plants of Mt Cameroon (Cable & Cheek 1998) and culminated in those for Dom (Cheek et al. 2010), Lebialem (Harvey et al. 2010) and Mefou in Central Region (Cheek et al. 2011). Two new national parks resulted from these checklists, the Mt Cameroon National Park, and the Bakossi Mts National Park, the first to be specifically for plant conservation (rather than for animals) in Cameroon.

Each of these conservation checklists has a section on Red Data species. Building on these, a national Red Data Book of the Plants of Cameroon was delivered (Onana & Cheek 2011), the first for any tropical African country. The Red Data book in turn helped the Tropical Important Plant Areas programme (Darbyshire et al. 2017) that resulted in the book on the Important Plant Areas of Cameroon that evidenced and mapped priority areas for plant conservation in Cameroon with local partners (Murphy et al. 2023).

In this paper we formally name a further new species to science from these surveys as *Tricalysia banyangmbo* Cheek & Sonké, allowing IUCN to accept a conservation assessment for the species, and for it then to be incorporated into conservation prioritisation initiatives.

## Material and methods

The methodology for the surveys in which this species was discovered is recorded in Cheek & Cable (1997). Nomenclatural changes were made according to the Code (Turland et al. 2018). Names of species and authors follow IPNI (continuously updated). Herbarium material was examined with a Leica Wild M8 dissecting binocular microscope fitted with an eyepiece graticule measuring in units of 0.025 mm at maximum magnification. The descriptions are based on herbarium specimens, spirit material and pictures when available, and data derived from field notes. The drawing was made with the same equipment with a Leica 308700 camera lucida attachment. Specimens were inspected from the following herbaria: BM, BR, K, P, WAG, YA using the conventions of Davies et al. (2023) and specimens were also viewed on GBIF (continuously updated), and species records checked on POWO (continuously updated). The format of the description follows those in other papers describing new species in Coffeeae e.g. Cheek et al. (2018a). Terminology follows Beentje & Cheek (2003), but for specialised structures of Rubiaceae, e.g. for stipule colleters, and domatia, generally follows Robbrecht (1987a, 1988). Phytogeographical considerations follow White (1979, 1983, 1993). All specimens cited have been seen unless indicated “n.v.” The conservation status of the new species was assessed according to IUCN Red List Categories and Criteria, version 3.1 (IUCN 2012; IUCN Standards and Petitions Subcommittee 2015).

The extent of occurrence and the area of occupancy were estimated with GeoCAT (http://geocat.kew.org) with a grid size of 2 × 2 km‥ Herbarium codes follow Index Herbariorum (Thiers continuously updated).

## Results

***Tricalysia banyangmbo*** Cheek & Sonké, **sp. nov**. Type: Cameroon, South West Region, Banyang Mbo Wildlife Sanctuary, Research station path to sanctuary via Nlowoa and Mbu river crossings, 5°20′ 14”N, 9°28′ 00”E, 200m alt., 24 Nov. 2000, fl., *Cheek* 10567 with Owens, Okon, Hinchcliffe, Rex, Suleiman, Myanzi, Kamwela & Johnston (holotype: K, barcode K000593391; isotypes: BR, SCA, YA). Fig 1 Evergreen shrub, 1.5 m tall, main axis (orthotropic stem) erect, dull white, subterete, c. 7 mm wide near base, internodes opposite and decussate 1.5 – 2.2 cm long; plagiotropic (axillary) stems 20 – 42 cm long, internodes at first dark brown to black, terete, 3 – 11 per stem, 1.25 – 1.8 mm diam., 2 – 3(– 5) pairs of leaves present per stem near apex, leaves subequal in shape and size; the longer stems sometimes with a pair of short side branches, indumentum of young stems (extending to the petioles, midribs, abaxial secondary nerves and abaxial stipule surfaces) of dense (c. 80 % of surface covered) erect, simple, translucent hairs 0.2 – 0.25 mm long, apex rounded, epidermis and indumentum persisting until the 3 – 5^th^ internode, decortication taking place gradually over several internodes, exposing a pale brown, glabrous, subglossy, lenticel-free surface. Stipules persistent, 2 – 2.5 × 1.75 – 2.75(– 3.5) mm, sheathing, limb hemi-orbicular, 1.25 – 1.5 mm long, awn apical to subapical, shorter than blade, 0.75 – 1 × 0.2 – 0.25(– 0.5) mm, outer surface indumentum as stem apex, inner surface pale yellow, glabrous apart from colleters, colleters winged type (Robbrecht 1987a), inserted towards base of stipule and adhering to the surface, 7 – 9 spaced equally along the width of the stipule, dorsiventrally flattened, dark brown, narrowly oblong, c. 0.75 × 0.2 mm, apex rounded Leaf-blades papery, upper surface drying dark grey-brown, lower surface greenish white, the midrib dark brown, secondary nerves orange-brown, elliptic or elliptic-oblong, less usually oblanceolate, (5.8 –)6 – 10.1(– 11.3) × (2.1 –)2.8 – 3.9 cm, apex acuminate, acumen (0.2)0.4 – 1.0(– 1.4) × 0.3 – 0.5 cm, base rounded, rarely minutely and abruptly cordate, abruptly slightly decurrent down the distal petiole; midrib sunken on adaxial surface, raised abaxially, camptodromous (secondary nerves not looping), domatia absent, secondary nerves (3 –)4(– 5) on each side, arising at 50 – 60 degrees from the midrib, curving slightly upwards and becoming parallel to the margin, midrib and secondary nerves on both surfaces with indumentum as stem but 30 – 40% cover, sparser and shorter on secondary nerves and leaf margin, otherwise glabrous. Tertiary nerves just visible with the naked eye, pale brown or dull yellow, scalariform, arising perpendicular to the midrib, parallel to each other, 10 – 15 between each secondary nerve, tertiary nerves sometimes bifurcating; quaternary nerves inconspicuous.Petiole distally canaliculate, proximally plano-convex, 2 – 4(– 5) × 1.2 – 1.5 mm, indumentum as stem. Inflorescences single (rarely a second superposed), in both axils of a node (often not developed in one axil), fertile nodes per stem 2 – 5, occurring at the 2 – 10 th node from the apex, flowers per inflorescence 1 – 3(– 5), inflorescence highly condensed; calyculi (2 –)3(– 4), cup-shaped, completely concealing the axis, 1^st^ order (proximal) calyculus brown, papery, shallowly cup-shaped, (0.2 –)0.75 – 0.8 × 1 – 2 mm, subtending 2^nd^ order calyculi, margin sometimes with short awns, surface with a few white adpressed straggling hairs 0.5 – 1 mm long. 2^nd^ order (medial) calyculus, 0.75 – 1 × 1.25 – 1.5 mm, sometimes absent, split, rarely frondose (leafy), the foliar lobes then two, narrowly oblong, c. 3 × 1 mm, stipular lobes if present, 1 – 2, triangular, minute, surface c. 80% covered in variously oriented simple white hairs 0.1 – 0.2 mm long, subtending 1 – 3 flowers, each with a single 3^rd^ order (distal) calyculus; sometimes with a more or less identical supplementary calyculus subtending additional flower(s); 3^rd^ order calyculi, tube (1 –)1.25 – 1.5 × 1.3 – 2 mm, entire or longitudinally split once (Fig. 1E&F), foliar lobes 2, subulate, (0.2 –)1.1 – 2.5 mm long, erect, resembling calyx lobes and decurrent as a rib down the calyculus tube, (rarely frondose), stipular lobes absent or inconspicuous, outer surface of tube and lobes densely covered (c. 80% of surface) in bright white adpressed simple hairs 0.1 – 0.2 mm long, inner surface densely white adpressed sericeous hairy, hairs 0.25 – 0.3 mm long, 100% cover, colleters not seen, pedicels absent or minute, < 0.1 mm long, concealed by calyculus. Flowers hermaphrodite, homostylous, 4 – (– 5)merous, sweetly scented of *Gardenia* at 4 pm (*Cheek et al*. 10567). Ovary-hypanthium 0.5 – 1 × 0.5 – 0.75 mm, calyx tube (0.75 –)1.5 – 2 × 1.25 – 1.5 mm, proximal part with ovary-hypanthium concealed within 3^rd^ order calyculus,; calyx lobes subulate, stout, erect, 1.45 – 3(– 3.5) × 0.15 – 0.25 mm, often varying in length in one flower by several 1/10ths of a mm, outer indumentum as distal calyculi 30 – 100% hair cover, hairs 0.2 – 0.75 mm long, inner surface densely hairy as in distal calyculi, colleters extremely inconspicuous and sparse, hidden among hairs, scattered, brown, 0.25 mm long. Corolla white, tube 3.5 – 4.5 × 0.4 – 0.8 mm, dilating to 2 mm wide at mouth, glabrous outside, apart from at the junction with the lobes, inside with hair band 1.6 – 1.9 mm deep, inserted 0.5 mm below mouth of tube, and c. 1 mm from base, hairs 0.2 – 0.25 mm long, adpressed, otherwise glabrous. Corolla lobes oblong, 4 – 7 × 1 – 1.5 mm, apex acuminate, often slightly laterally compressed (Fig. 1G), adaxially glabrous, abaxially white sericeously hairy, hairs 0.1 – 0.12 mm long, ±adpressed, 50 – 80% cover on middle and distal part, extending to the margins reducing to 40% on proximal part and extending to the apex of the tube (Fig. 1E), Stamens almost fully exserted, glabrous, anthers oblong, 2 – 3.6 × 0.3 – 0.4 mm, submedifixed, affixed 1/4– 1/3 from the base, apical connective appendage absent or vestigial; filaments 1.7 – 1.75 mm long, inserted just below mouth of tube, dorsiventrally flattened, c. 0.25 mm wide at base, tapering to 0.1 mm wide at apex. Disc subcylindrical, apex concave, 0.2 – 0.3 × 0.4 mm, glabrous Style 5 – 6 × 0.15 – 0.25 mm (narrowest at base, widest at apex), exserted, basal 2 mm glabrous, above moderately densely hairy with adpressed to spreading hairs 0.1 – 0.12 mm long; style head 0.8 mm long, glabrous, papillate, diverging into two recurved stigmatic lobes 1.2 – 1.3(– 2) mm long, apices acute, adaxial surfaces papillate. Ovary 2–locular, ovules 2 per locule, superposed, immersed in the swollen placentae. Fruit and seed unknown.

**Figure 1.**
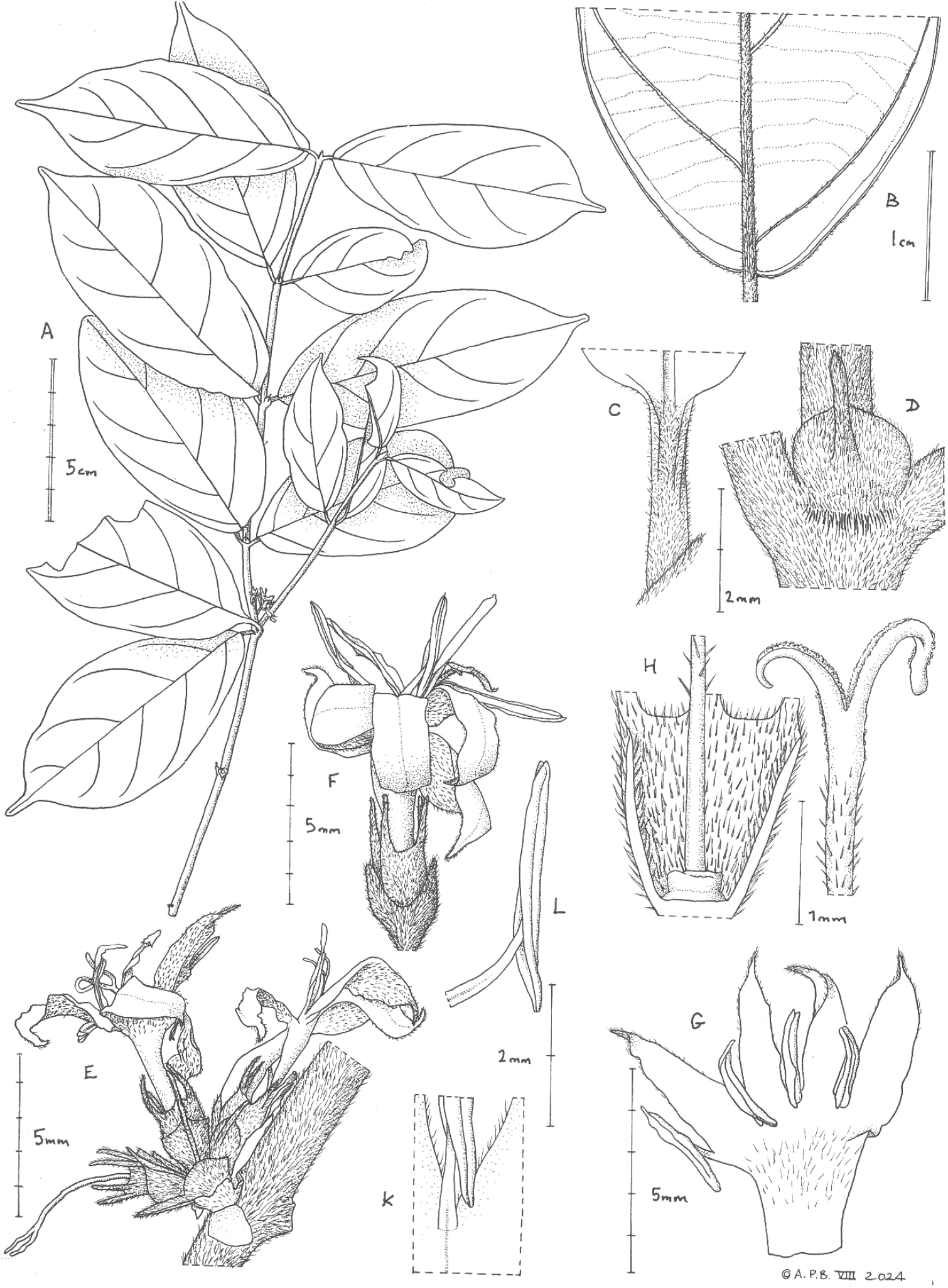
Tricalysia banyangmbo. **A**. Habit, flowering stem. **B**. Lower part of leaf, abaxial surface. **C**. Petiole, adaxial surface. **D**. Stipule, *in situ*. **E**. Inflorescence (atypical 5-merous flowers). **F**. Flower, side view, hydrated. **G**. Corolla opened to show inner surface. **H**. Calyx tube sectioned to show disc and style base. **J**. Stigma. **K**. Staminal filament *in situ* on corolla inner surface. **L** Stamen, side view. A–F & H–L from *Cheek* 10564 (K); G from *Sonké* 2411 (K), all drawn by Andrew Brown.

**Map1.**
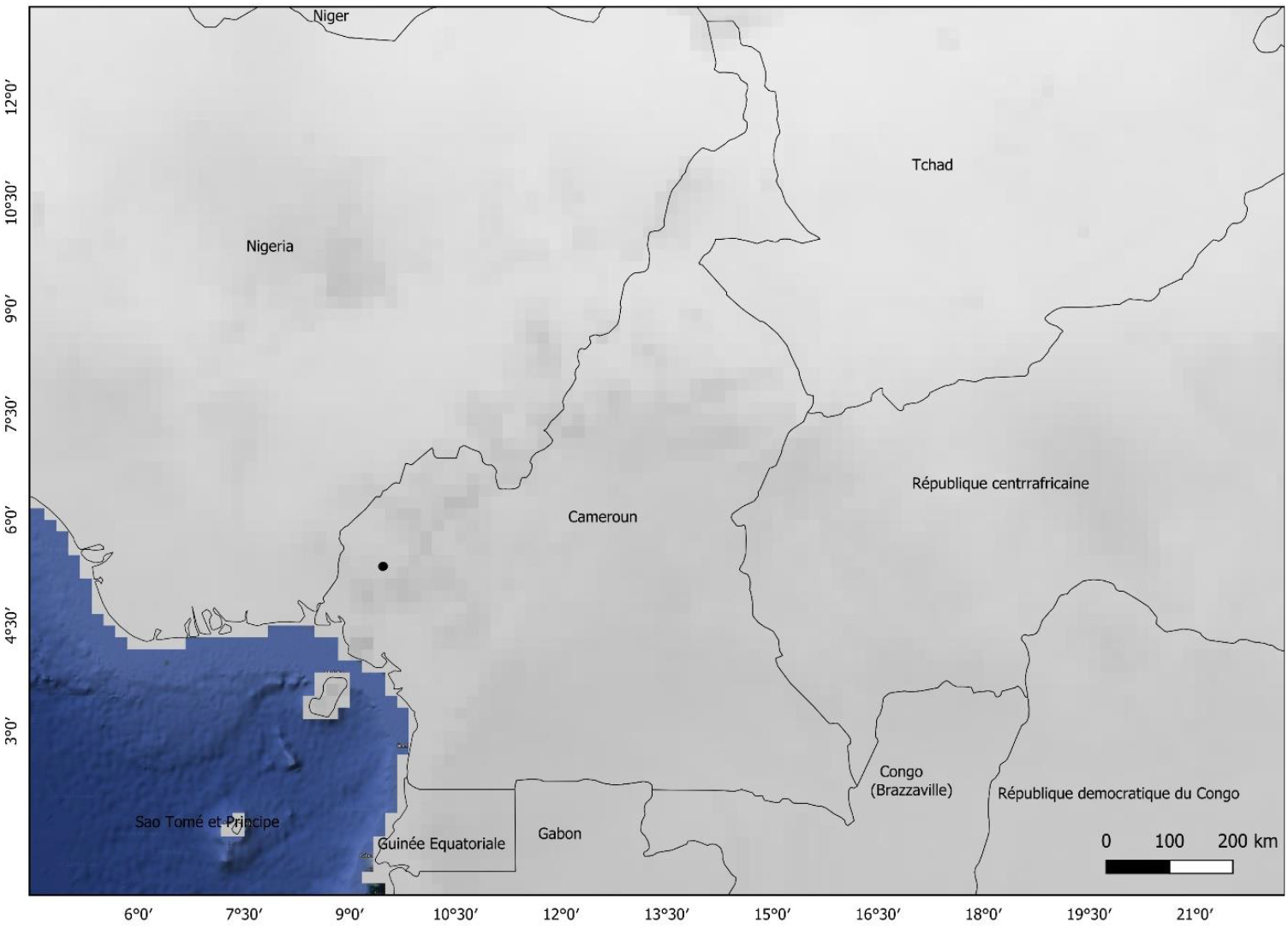
Global distribution of *Tricalysia banyangmbo*

## RECOGNITION

Similar to *Tricalysia sylvae* Robbr. of Sect. *Tricalysia* in the subulate hairy calyx lobes as long as the calyx tube or longer, in the condensed inflorescences, and the dense erect indumentum of the young stems, petioles and midribs, the hairs extending to the style and exterior corolla lobes (but not the anthers), differing in the flowers 4(– 5) merous (vs (5 –)6-merous), the stipule blade hemiorbicular, longer than the 0.8 – 1 mm long awn (vs triangular, awn longer than blade), the leaves camptodromous (vs brochidodromous), anther apex obtuse, lacking appendage (vs with connective appendage), domatia absent (vs present) (Table 1).

## DISTRIBUTION & ECOLOGY

Lower Guinea subcentre of endemism (White, 1979). This species occurs in Cameroon, known only from the vicinity of Banyang Mbo Wildlife Sanctuary near Nguti, SW Region, understorey shrub of lowland evergreen forest; c.200 m alt. Map 1.

## SPECIMENS EXAMINED CAMEROON

**South West Region, Banyang Mbo Wildlife Sanctuary**, Research station path to sanctuary via Nlowoa and Mbu river crossings, 5°20′ 14”N, 9°28′ 00”E, 200m alt., fl., 24 Nov. 2000, *Cheek* 10567 with Owens, Okon, Hinchcliffe, Rex, Suleiman, Myanzi, Kamwela & Johnston (K holo., barcode K000593391; isotypes: BR, MO, SCA, YA); 3 km after the research station of WCS, between the rivers Nloa & Mbu, 5°20′ 12”N, 9°28′ 00”E, 200m alt., fl., 27 Nov. 2000, *Sonké* 2373 with Zapfack, Sainge, Ngwa, Okon (BR, K, MO, SCA, YA); 4 km after the research station of WCS, after the river Mbu, 5°20′ 20”N, 9°28′ 40”E, 200 m alt., fl., 28 Nov. 2000, *Sonké* 2411 with Pollard, Sainge, Okon (BR, K, SCA, YA).

## CONSERVATION ASSESSMENT

*Tricalysia banyangmbo* is known from three collection sites which are all in close proximity to each other, the most distant c. 1 km apart from the others, equating to one threat-based location on the southern edge of the Banyang Mbo Wildlife Sanctuary (BMWS). The species is also threatened by the low number of individuals recorded: no more than three have been seen (the authors, pers.obs.). Using the IUCN preferred 4 km^2^ grid cells, the extent of occurrence is 4 km^2^ and the area of occupancy is similar. *Tricalysia banyangmbo* on the evidence available, is highly localized, being known only from three individuals collected at the southern edge of the Banyang Mbo Wildlife Sanctuary. This is despite the fact that a team of ten plant collectors, with 20 support workers, spent two weeks collecting specimens in the area in November and December 2000 (Cheek 2001).

Moreover, the two authors have made several lengthy visits to the Banyang Mbo forest some years ago without locating this taxon. *Tricalysia banyangmbo* is unknown from the Korup National Park (to the west) or from the Bakossi Mts (to the south). Both of these areas have been subject to botanical inventory, although not exhaustively so. Further botanical surveys in subsequent years from the same base near Nguti by botanists from the University of Rostok, Germany led by Prof. Stefan Porembski (pers. comm. to Cheek) and the second author of this paper (BS of Higher Teachers’ Training College, University of Yaoundé I) have similarly not produced any additional records of the species so far as is known. Neither has this species been found elsewhere in botanical surveys in SW Region Cameroon and surrounding areas (Cheek et al. 1992; Cheek *et al*. 1996; Cable & Cheek 1998; Cheek *et al*. 2000; Maisels *et al*. 2000; Chapman & Chapman 2001; Cheek *et al*. 2004; Harvey *et al*. 2004; Cheek *et al*. 2006; Cheek *et al*. 2010; Harvey *et al*. 2010; Cheek *et al*. 2011).

The only known location for *Tricalysia banyangmbo* is within the boundary of the area proposed as an oil palm plantation by the U.S.A.-based company, Herakles (http://www.greenpeace.org/usa/wp-content/uploads/legacy/Global/usa/planet3/PDFs/HeraklesCrimeFile.pdf. Although this operation was reported as suspended in 2013 there have been concerns that it will be resurrected in the future since Herakles subsequently continued its activities on the ground (https://www.grain.org/article/entries/5037-communities-lose-out-to-oil-palm-plantations, downloaded 19 Sept. 2024). Due to insecurity in the English-speaking parts of Cameroon since 2016, resulting in many deaths and the displacement of tens of thousands of many citizens, plans for industrial operations in the region that includes Banyang Mbo have had to be suspended. However, as fighting appears now to be on the decrease it is likely that these will resume including the oil palm plantation plan that will impact Banyang Mbo.

Therefore, there is every chance in the future that all those individuals of the species known will become extinct when clearing proceeds for the plantation.

Even if a broad buffer area were created outside the boundary of Banyang Mbo, there is a risk that farmers displaced by the future plantation project will take up land within it and so threaten the only known locality of the species. For these reasons we consider the only known location for the species to be highly threatened and therefore assess *Tricalysia banyangmbo* as Critically Endangered CR B1ab(iii) + B2ab(iii).

It is possible that the species also extends to other areas nearby the BMWS where it might have the prospect of protection. Yet areas adjacent e.g. South and west of Nguti continue into the main part of the area designated for plantation by Herakles. This area remains relatively unsurveyed for plants, although it occurs within the area of Tropical Africa with the highest recorded density of wild plant species, which occurs in SW Region Cameroon (Barthlott et al. 1996)‥

While BMWS is protected mainly for its animals, it has also been designated as an Important Plant Area (CMNTIPA030, Murphy et al. 2023), as has the neighbouring area to the south and west of Nguti, as the Nguti Forests (CMNTIPA043, Murphy et al. 2023).

## ETYMOLOGY

Named (indeclinable word in apposition; art. 23.1 and 23.2 of the code, Turland et al. 2018) for the Banyang Mbo Wildlife Sanctuary to which this species on current knowledge is unique.

## NOTES

The new species, *Tricalysia banyangmbo*, shares so many features with *T. sylvae*, differing only quantitively for the main part, that shared recent common ancestry seems highly probable. However phylogenetic study is needed to test this hypothesis. Were it not for the few additional qualitative characters that separate the taxa, subspecific not specific status would have been opted for. These two lowland forest taxa are separated geographically by the Cameroon Highlands, *T. sylvae* to the east where it appears relatively common, and *T. banyangmbo* to the west. Although some lowland areas occur within the highlands between the ranges of the two species e.g. the Chide valley, neither species has been detected there, despite quite high botanical survey effort (Cheek et al. 2004). It is possible that at one time that there was a single widespread species that became separated into two as the Highlands developed. Equally it is possible that one taxon arose from the other by a dispersal event: the seeds are carried in fleshy red fruits and so are probably bird-dispersed.

A remarkable feature of both species is that the subulate calyx lobes exceed the calyx tube in length, a feature seen in only five other species of the Section. None of the other five species share the densely patent hairy indumentum of the young stems.

Some variation was noted within the species. While *Sonké* 2373, 2411 are 4-merous, several 5-merous flowers were found in *Cheek* 10567.

## Discussion

### The endemic and threatened plant species of Banyang Mbo Wildlife Sanctuary (BMWS)

The BMWS covers an area of about 662 km^2^, between 05°08’ and 05°34’ N and 09°30’ and 09°47 E. It spans an altitudinal range of c.120 – 1760m. The mean annual rainfall is 4082.7±486mm with heaviest rains in July-September. The mean annual maximum temperature is 30.2ºC and the minimum 23.7ºC (Nchanji and Plumptre 2003; Murphy et al. 2023). Apart from *Tricalysia banyangmbo*, several other species are endemic or near endemic to Banyang Mbo Wildlife Sanctuary or otherwise highly range-restricted and threatened. These can be divided into two groups based on habitat.

### A.Lowland evergreen forest

Growing in the same habitat and general area as *Tricalysia banyangmbo*, are *Tricalysia lejolyana* Sonké & Cheek (EN, Rubiaceae, Sonké et al. 2002a) and *Warneckea ngutiensis* R.D.Stone (CR, Melastomataceae-Olisbeoideae, Stone & Cheek 2018), both also endemic and collected in the same survey period in 2000 as *Tricalysia banyangmbo*. Other new species to science resulting from this survey and found in the same vicinity in lowland forest were *Cola metallica* Cheek (CR, Malvaceae Sterculioideae, Cheek 2002 also found e.g. in the Bakossi and Mt Cameroon areas), *Psychotria bimbiensis* Bridson & Cheek (CR, Rubiaceae, Cheek & Bridson 2002, also in Bimbia-Bonadikombe near Limbe). Subsequently, the area was also found to contain the largest global population of the threatened orchid *Ossiculum* P.J.Cribb & Laan (EN, now *Calyptrochilum aurantiacum* (P.J.Cribb & Laan) Stévart, M.Simo & Droissart (Simo-Droissart et al. 2018) which formerly had been considered possibly globally extinct (Onana & Cheek 2011). Several highly threatened species previously known from very few other locations were also found, e.g. *Allophylus conraui* Gilg ex Radlk. (VU, Sapindaceae, Cheek & Etuge 2009b), *Ancistrocladus grandiflorus* Cheek (VU, Ancistrocladaceae, Cheek 2002) and *Vepris letouzeyi* Onana (EN, Rutaceae, Onana & Chevillotte 2015).

### B.Submontane (cloud) forest

This habitat occurs in the east and NE of Banyang Mbo. New species to science discovered in this habitat at BMWS were *Aulacocalyx mapiana* Sonké & Bridson (VU, Rubiaceae, Sonké & Bridson 2001, endemic to Banyang Mbo. Ngovayang and Rumpi Hills), *Rothmannia ebamutensis* Sonké (EN, Rubiaceae, Sonké 2000, endemic to BMWS) and *Rinorea fausteana* Achoundong (EN, Violaceae, Achoundong & Cheek 2003, also known from Rumpi Hills and Bakossi Mts and *Manniella cyprepedioides* Salazar et al. (EN, Orchidaceae, Salazar et al. 2002 also known in Bakossi and a few other locations). Sixteen other extremely rare orchid species have been found in BMWS, mainly in submontane forest, such as *Polystachya kornasiana* Szlach. & Olsz. (mainly EN, Orchidaceae, Droissart et al. 2006). To these can be added *Rhipidoglossum paucifolium* D.Johanss. (Lachenaud et al. 2013), likely EN although not yet listed on iucnredlist.org.

Some submontane species thought endemic to the Bakossi Mts immediately to the south, were also unsurprisingly found in BMWS on the 2000 survey, namely *Octoknema bakossiensis* Gosline & Malécot (EN, Gosline & Malécot 2011), *Coffea bakossi* Cheek & Bridson (EN, Rubiaceae, Cheek et al. 2002a), while others were not (e.g. *Coffea montekupensis* Stoff. (VU, Rubiaceae, Stoffelen et al.1997) even though this species extends far beyond BMWS to e.g. Lebialem (Harvey et al. 2010). Meanwhile, *Salacia conraui* Loes. (CR, Celastraceae, Loesener, 1910), historically recorded from the Ntale Mts of BMWS, has not been refound in more than 125 years and is listed as potentially extinct (Onana in Murphy et al. 2023: 18).

In total five globally Critically Endangered, 17 Endangered and 21 Vulnerable plant species are listed for BMWS (iucnredlist.org; Murphy et al. 2023). This number is likely to grow if botanical survey work continues, since only a small fraction of the area of the BMWS, perhaps 20%, has been surveyed, these parts being in the south while the remainder is botanically unknown. Apart from new species to science having been discovered, new chemical compounds to science have also been found. Five new undescribed *ent-*abietane diterpenoids from the rare species *Suregada occidentalis* (Hoyle) Crozat (Euphorbiaceae) in Banyang Mbo in have been named in honour of Banyang Mbo as banyangmbolides A-E. These may have future applications for humanity (Olaranont et al. 2024). Clearly BMWS is globally important for conservation for plants, despite being incompletely surveyed.

## Conclusion

Cameroon has the highest number of globally extinct plant species of all countries in continental tropical Africa (Humphreys et al. 2019). The extinction of species such as *Oxygyne triandra* Schltr. (Thismiaceae, Cheek et al. 2018b) and *Afrothisia pachyantha* Schltr. (Afrothismiaceae, Cheek & Williams 1999; Cheek et al. 2019a; Cheek et al. 2023a) are well known examples, recently joined by species such as *Vepris bali* Cheek (Rutaceae, Cheek et al. 2018c) and *Vepris montisbambutensis* Onana (Onana & Chevillotte 2015). However, another 127 potentially globally extinct Cameroon species are documented (Murphy et al. 2023: 18 – 22).

It is critical now to detect, delimit and formally name species as new to science, since until they are scientifically accepted they are essentially invisible to science. Only when they have a scientific name can their inclusion on the IUCN Red List be facilitated (Cheek et al. 2020b). Most (77%) species named as new to science in 2020 were already threatened with extinction (Brown et al. 2023). Many new species to science have evaded detection until today because they have minute ranges which have remained unsurveyed, as was the case with *Tricalysia banyangmbo*.

If further global extinction of plant species is to be avoided, effective conservation prioritisation, backed up by investment in protection of habitat, ideally through reinforcement and support for local communities who often effectively own and manage the areas concerned, is crucial. Important Plant Areas (IPAs) programmes, often known in the tropics as TIPAs (Darbyshire et al. 2017; Murphy et al. 2023) offer the means to prioritise areas for conservation based on the inclusion of highly threatened plant species, among other criteria. Such measures are vital if further species extinctions are to be avoided of narrowly endemic, highly localised species such as *Tricalysia banyangmbo*.

## Acknowledgements

We thank two anonymous reviewers for reviewing an earlier version of this manuscript. The botanical surveys in Cameroon which resulted in this paper were mainly supported by Earthwatch Europe (1993 – 2005) and by the Darwin Initiative of the UK Government through the Plant Conservation of Western Cameroon, and the Red Data Book of Cameroon projects, both led by RBG, Kew, working with the IRAD-National Herbarium of Cameroon. Gaston Achoundong, former head of the National Herbarium of Cameroon (YA) and his successors including Jean Michel Onana and Barthelemy Tchiengué are thanked for their collaboration and support over the years. Our thanks also go to Jeannette Mapi-Sonké and the authorities of the University of Yaoundé I for their support to the second author’s research activities.

## Declarations

